# What can a neuron compute

**DOI:** 10.64898/2026.06.08.730984

**Authors:** Ido Aizenbud, David Beniaguev, Noam Pnueli, Idan Segev, Michael London

## Abstract

Cortical pyramidal neurons possess elaborate dendritic trees with diverse nonlinear membrane conductances and thousands of plastic synapses, suggesting substantial computational capabilities at the single-cell level. Yet, what can a neuron compute remains an open question, largely due to the lack of a systematic framework to quantify its computational capabilities. We introduce *TwinProp*, a digital-twin-based backpropagation algorithm that enables gradient-based optimization of synaptic strengths and dendritic locations in detailed neuron models via a millisecond-accurate deep neural network (DNN). Using *TwinProp*, we demonstrate that a detailed model of rat layer 5 pyramidal cell (L5PC) can perform naturalistic image and audio classification tasks at a remarkably high accuracy, significantly surpassing perceptron and leaky integrate-and-fire baselines. The same neuron solves high-dimensional nonlinear problems, including exclusive-or (XOR), 10-bit parity, and random Boolean tasks, demonstrating capabilities typically attributed to multilayer networks. Mechanistically, increasing task complexity recruits distributed dendritic nonlinearities, including NMDA- and voltage-dependent mechanisms; removing these or collapsing dendritic structure markedly impairs performance. These findings identify dendrites as a substrate for high-order feature binding and position single cortical pyramidal neurons as powerful, noise-robust, general-purpose analog computational units. Our results offer testable in vivo predictions and provide a systematic framework linking cellular morpho-electrical properties to computation in both brains and artificial systems.

## Introduction

In view of recent advances in artificial intelligence, modern neuroscience faces a fundamental paradox. On one hand, artificial Deep Neural Networks (DNNs), composed of neuron-inspired simple computational units, not only rival human performance on various complex tasks but also develop internal representations partially aligned with neural activity, e.g., in the primate visual ventral stream (Yamins and DiCarlo 2016; Khaligh-Razavi and Kriegeskorte 2014; Yamins et al. 2014). This success has been taken to suggest that the computations performed by individual neurons can be safely abstracted away in favor of the collective computations of vast networks composed of simple units (McCulloch and Pitts 2021; LeCun et al. 2015; Sejnowski 2020).

By contrast, decades of increasingly refined anatomical and physiological studies have revealed that biological neurons are starkly different from the simplified units used in present-day AI systems. Cortical pyramidal neurons receive thousands of synaptic inputs distributed over highly-branched dendritic trees, where inputs interact nonlinearly at many separate locations before generating axonal output spikes (Stuart and Sakmann 1994; Larkum et al. 1999; Gidon et al. 2020; Polsky et al. 2004; Magee 2000; Golding et al. 2002). The resulting input-output mapping is governed by a rich set of factors: passive cable properties, dendritic membrane nonlinearities, overall dendritic morphology (Spruston 2008), and the location and strength of synapses, which are themselves continually reshaped by learning (Malenka and Bear 2004; Lamprecht and LeDoux 2004; Citri and Malenka 2008).

This biological reality suggests that individual neurons may act as powerful computational devices in their own right (Segev and Rall 1998; London and Häusser 2005). However, the extent of single neuron computational capabilities remains unknown, and without this knowledge, the cost of abstracting single-neuron complexity away cannot be assessed. A critical question therefore arises: what can a single, morphologically and electrically complex neuron compute? Concretely, as task complexity increases, does there exist a configuration of synaptic strengths and dendritic locations that enables the neuron to solve the task? Because current experimental methods cannot comprehensively access or manipulate all synaptic inputs in biological tissues, this question can presently be addressed only in biologically realistic simulations. Importantly, we focus here on what can neurons compute, not on how such synaptic configurations are learned.

Exploring this question has proven historically difficult. Early abstractions, including the perceptron (Rosenblatt 1958) and the leaky integrate-and-fire model (Abbott 1999), reduced neuronal computation to linear summation and thresholding, which fundamentally limits expressivity (Cover 1965; Torres-Moreno et al. 2002; Minsky and Papert 2017). Subsequent pioneering theoretical work established that pyramidal neurons can be approximated as two-layer computational units (Poirazi and Mel 2001; Poirazi et al. 2003). Abstract follow-up models showed that such two-layer structures can solve non-trivial tasks (Jones and Kording 2021; Agrawal and Buice 2025), but these approaches omit spiking dynamics and active dendritic mechanisms. Conversely, detailed compartmental models capture biological realism (Rall 1957, 1959, 1960, 1967; Segev and Rall 1988; Koch and Segev 2000; Segev and London 2000) yet are analytically intractable, and have traditionally been explored through manual parameter tuning (Mel 1992). As a result, although detailed neurons can match perceptron-like classification performance (Moldwin and Segev 2020), attempts to solve nonlinear problems like XOR have yielded sub-optimal results (Bicknell and Häusser 2021; Deistler et al. 2024).

This raises a fundamental question: are these limitations intrinsic to detailed neuron models, or an artifact of the optimization methods used to probe them? Addressing this is challenging, as optimizing synaptic strengths and dendritic locations in compartmental models is computationally costly, and spike generation introduces non-smooth dynamics that hinder gradient-based optimization. Consequently, a systematic framework for jointly optimizing functional (strength) and structural (dendritic location) synaptic weights remains lacking.

Here we introduce *TwinProp*, a digital-twin-based backpropagation algorithm that addresses this challenge. Leveraging a differentiable DNN twin of the neuron’s input–output function (Beniaguev et al. 2021; Beniaguev 2023), *TwinProp* enables efficient optimization of both synaptic strengths and dendritic locations in detailed neuron models. We show that a single layer 5 cortical pyramidal neuron can implement complex nonlinear computations, from naturalistic sensory classification to high-dimensional abstract problems. We further identify the morphological and physiological dendritic mechanisms that support these computations and provide a principled route for incorporating dendritic computation into circuit-level network models.

## Results

### Challenging a single neuron with complex tasks using the *TwinProp* algorithm

Figure 1 outlines the steps taken to apply the *TwinProp* algorithm, which casts the question *“what can a single neuron compute?”* as an optimization problem over synaptic weights. The objective is to find, via gradient-based optimization, a configuration of synaptic strengths and dendritic locations that enables a neuron to solve a given computational task. For a given task, for example, classifying images of cats versus dogs (Fig. 1a, top left), inputs are first transformed into biologically plausible population spike trains using an encoder (Fig. 1a, green and red rasters). The neuron processes these spatiotemporal inputs, and its output spike train is mapped to a task label (e.g., “cat” vs. “dog”) by a decoder within a predefined decision window (Fig. 1a, right). Successful classification requires generating at least one spike for target stimuli (e.g., “cats”) and no spike for non-target stimuli (e.g., “dogs”).

**Figure 1.**
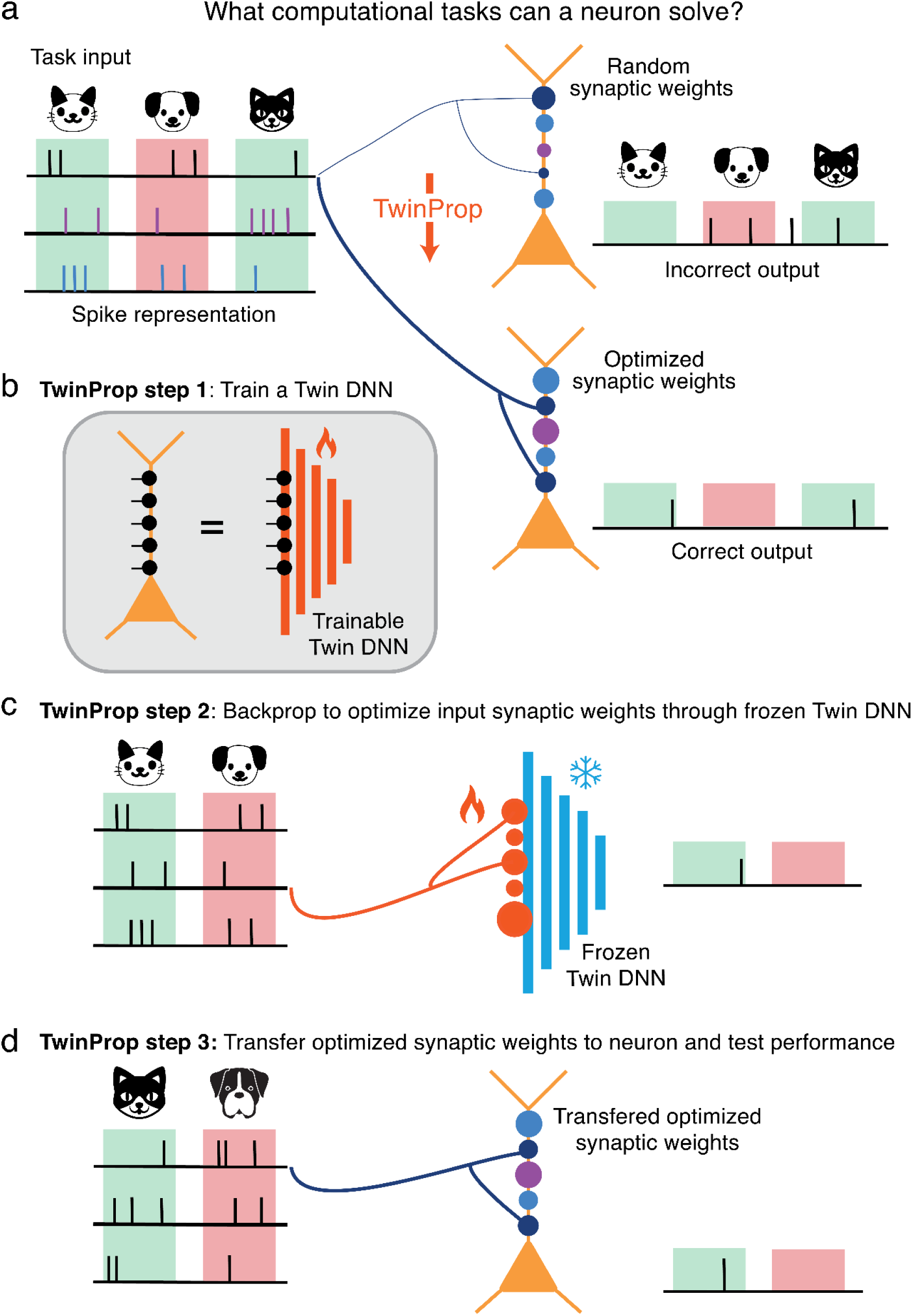
*TwinProp* algorithm for probing the range of tasks that a single neuron can implement. **a.** Schematic exposition of the algorithm: given a computational task, e.g., an image classification task (top left), it is first encoded into a biologically-plausible spiking representation (green/red boxes). The neuron model needs to solve the task based on these input spikes. Top right: with random synaptic weights (random synaptic strengths and locations), the neuron fails to solve the task. Bottom right: the *TwinProp* algorithm finds a set of optimized weights such that the neuron solves the task (spikes for cats; no spikes for dogs). **b-d**. Steps in applying *TwinProp* algorithm. **b**. step 1: training a digital twin DNN (right) to match the I/O of the detailed model (left) under a large set of random inputs and synaptic weights. **c.** step 2: freezing the internal parameters of the twin DNN (blue DNN) and optimizing only the weights of the input synapses (orange) such that the DNN solves the task. **d.** step 3: mapping the optimized synaptic weights from the DNN twin back onto the detailed neuron model and verifying that it correctly solves the task.

With random synaptic strengths and dendritic locations, performance is at chance level (Fig. 1a, top right). The fundamental challenge is to identify synaptic configurations that maximize task performance. While this optimization is tractable for linear models with analytically characterized capacities, it remains a major challenge for detailed neuron models (Rall 1957, 1959, 1960, 1967; Segev and Rall 1988; Koch and Segev 2000; Segev and London 2000), for which no analytical characterization exists and one must rely on empirical search over a high-dimensional parameter space. The *TwinProp* algorithm developed here (Fig. 1a, bottom center) addresses this challenge by efficiently identifying synaptic weights that enable the neuron to solve the task (Fig. 1a, bottom right), if such a solution exists.

Beniaguev et al. (2021) demonstrated that the I/O relationship of a detailed layer-5 cortical pyramidal cell (L5PC) model can be accurately approximated by a deep neural network (DNN). The size of the required DNN underscored the complexity of this I/O relationship, while the approximation’s accuracy established the DNN as a differentiable digital twin for the detailed model at millisecond temporal resolution. *TwinProp* leverages this finding: because the DNN twin is differentiable, one can use backpropagation through it to compute gradients for the synaptic weights of the detailed model, optimizing them for a specific computational task (schematically depicted in Fig. 1b–d).

The *TwinProp* algorithm consists of three steps. First, we fit a DNN twin for the detailed model (Fig. 1b, Fig. S1). Briefly, we generate a large training dataset by driving the detailed neuron model with random input spike trains and random synaptic weights (varying both strength and dendritic location) and recording the output voltage and spikes for each case. We then use this dataset to train a DNN to reproduce the I/O of the neuron model for any given input, and validate its accuracy on held-out inputs and synaptic weights, verifying that it faithfully reproduces the somatic voltage as well as spiking output (Fig. S2). Second, we freeze the DNN’s internal parameters and optimize only the synaptic weights to solve the task. This is achieved by backpropagating gradients through the frozen network to minimize the task loss (Fig. 1c). Finally, we map the optimized synaptic weights (both strengths and locations) back to the detailed neuron model and test its performance (Fig. 1d). See **Methods** for a comprehensive description of the *TwinProp* algorithm.

The *TwinProp* algorithm allows us to systematically explore the space of computations that a single neuron could implement. Below we use the *TwinProp* algorithm to quantify the computational capabilities of a L5PC on a large set of tasks and uncover the dendritic mechanisms that enable these computations.

### A single L5PC solves naturalistic sensory classification tasks

We first ask whether a single L5PC can perform naturalistic computations from realistic spike trains that encode sensory input. To ensure the neuron received biologically plausible inputs rather than raw pixels or audio waveforms, images (Choi et al. 2020) and sounds (Warden 2018) were transformed into population-spike representations using canonical early sensory encoders (Fig. 2a1, a2), retina → LGN→ V1 → V2 (Riesenhuber and Poggio 1999), and cochlea → VCN (Pfeiffer 1966; Cramer et al. 2019), see **Methods** for full description of these encoders. These spike trains were mapped as distinct excitatory and inhibitory inputs onto a detailed model of rat L5PC (Fig. 2a1, a2). The connectivity was constrained to strictly conserve Dale’s law, while allowing multiple synaptic contacts per afferent axons and synaptic strengths bounded within realistic conductance ranges (Markram et al. 2015) (see **Methods** and **Fig. S3** for detailed connectivity parameters and weight distributions).

**Figure 2.**
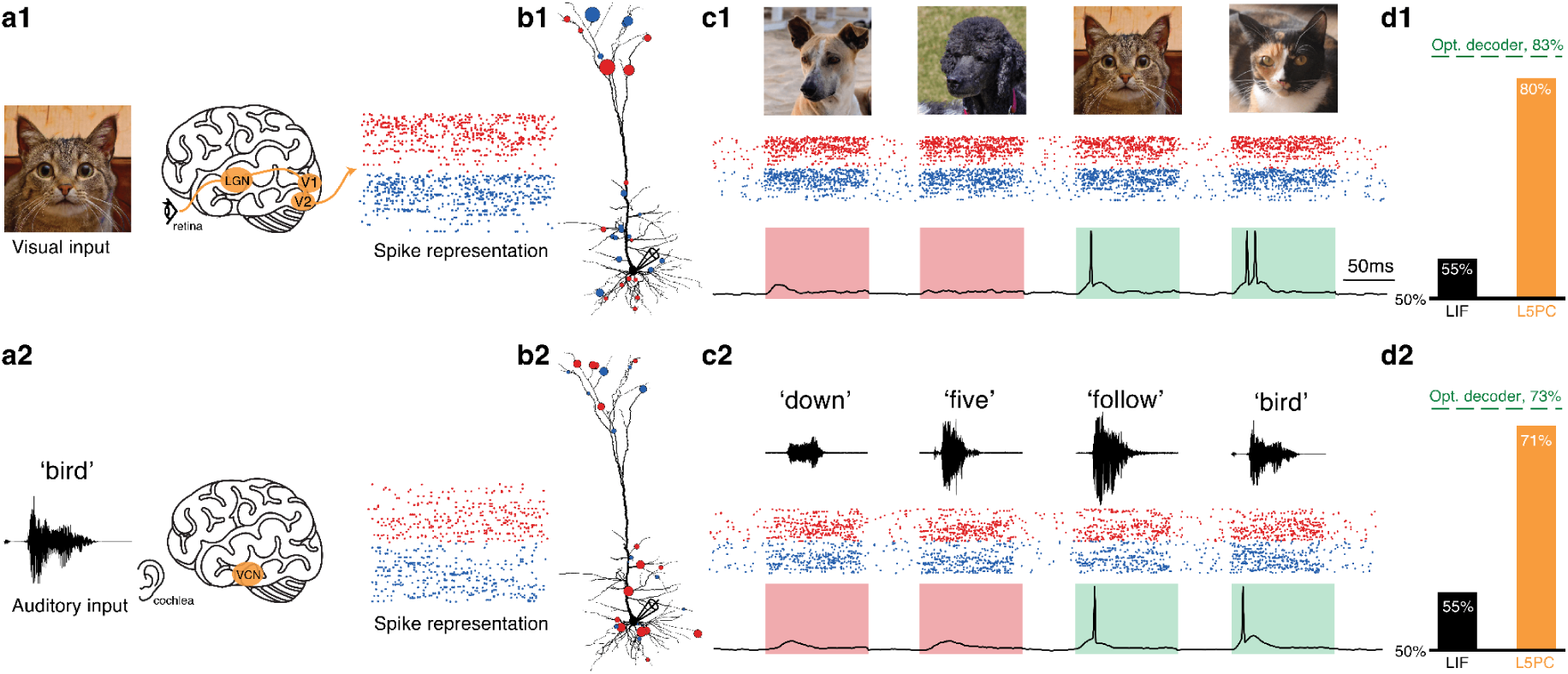
A single L5PC solves naturalistic visual and auditory classification tasks. **a1.** Visual pathway cartoon: image → retina → LGN → V1 → V2 like process produces a population spike representation of the input image (raster plot shows subset of excitatory and inhibitory input spikes in red and blue, respectively). **b1.** Modeled L5PC with optimized synaptic weights obtained by *TwinProp* for the visual task (top 5% strongest synapses shown; red – excitation, blue – inhibition). **c1.** Example test input images (cats/dogs) used (top) with their respective raster plots (middle, subset of excitatory and inhibitory spikes in red and blue, respectively) and the resultant somatic voltage response within the 100 ms decision windows (bottom; green = target class). **d1.** Test accuracy on the visual task (cats vs. dogs): L5PC ≈ 80%, approaching a near-perfect decoder (DNN upper bound, ∼83%), exceeding the LIF (∼55%) baseline. **a2.** Auditory pathway cartoon: auditory waveform → cochlea → ventral cochlear nucleus (VCN) like process produces a spike representation of the auditory input. Ten spoken words from the Speech Commands dataset were randomly assigned to two classes, creating an arbitrary binary classification task. **b2.** Modeled L5PC with optimized synaptic weights obtained by *TwinProp* for the auditory task (top 5% strongest synapses shown; red – excitation, blue – inhibition). **c2.** Example test input words (left; “down,” “five,” “follow,” “bird”) with respective raster plots (middle, subset of excitatory and inhibitory spikes in red and blue, respectively) and resultant somatic voltage traces (bottom). Here, both rasters and decision windows are 100 ms long. **d2.** Test accuracy on the auditory task (arbitrary two-class word classification: ‘backward’, ‘bird’, ‘dog’, ‘eight’, ‘follow’ speech commands, vs. ‘bed’, ‘cat’, ‘down’, ‘five’, ‘forward’ speech commands): L5PC ≈ 71% approaching near-perfect decoder (DNN upper bound, ∼73%), outperforming the LIF (∼55%) baseline.

The neuron’s goal was to produce at least one spike within a decision window for the positive classes (e.g., cats), but not for the negative classes (e.g., dogs). Using *TwinProp*, we optimized the neuron’s synaptic weights on a training set (resulting weight distributions, strengths and locations, are shown in Fig. S3 A1-F1, A2-F2) and evaluated generalization performance on a held-out test set (examples are shown in Fig. 2b1 and b2). To contextualize the single neuron’s performance, we compared it against two linear neuron models (perceptron and the leaky integrate-and-fire, LIF), each possessing the same number of synaptic weights as the L5PC. These baselines reflect what can be achieved by a point neuron that integrates inputs linearly; failure of these models to solve the task would indicate that it is not linearly separable. As a complementary reference, a high-capacity deep neural network (DNN) decoder (a fully connected neural network of up to 7 layers, see **Methods**) operating on the same spike-based inputs serves as an approximate “ideal observer,” providing an empirical upper bound on the performance attainable from the input spikes irrespective of biological constraints. Thus, if the L5PC’s performance approaches that of the DNN’s, it means that it effectively extracts most of the task-relevant information available in its inputs.

On the visual task of discriminating between pictures of cats and dogs (Choi et al. 2020), the single L5PC achieved ∼80% accuracy, exceeding linear perceptron (∼67%) and linear integrate-and-fire (∼55%) baselines that used the same spike features and approaching the performance of a DNN upper bound (∼83%) (Fig. 2c1, d1). To test generalization across sensory modalities, we next applied our framework to an auditory discrimination task using spoken commands recorded from multiple speakers (Warden 2018; Cramer et al. 2019). The neuron was required to spike selectively for an arbitrarily chosen set of five target commands (‘backward’, ‘bird’, ‘dog’, ‘eight’, ‘follow’) while remaining silent for the other five non-target commands (‘bed’, ‘cat’, ‘down’, ‘five’, ‘forward’). On this task, the same L5PC reached ∼71%, again surpassing the linear (∼60%) and LIF (∼55%) baselines and nearing the DNN upper bound (∼73%) (Fig. 2c2, d2). Example trials (depicted in Fig. 2) show reliable decision-window spiking aligned with class labels, indicating a consistent single-cell readout from distributed sensory spikes.

Together, these results demonstrate that spatiotemporal dendritic integration at the single-cell level can implement classification on complex sensory inputs. For the sensory tasks we probed, the performance of the L5PC was very close to that of the DNN “ideal observer” upper bound, which itself does not reach 100% accuracy because the spike-based encoders and the finite, noisy datasets do not preserve perfectly separable class information. However, while these results demonstrate competence on complex naturalistic tasks, the encoding involves hundreds of binary inputs (see **Methods**), making it difficult to characterize the precise function the neuron is computing. We therefore next ask whether the L5PC could solve tasks defined over a more compact, abstract feature space, where the computational complexity can be systematically manipulated.

### A single L5PC solves high-dimensional Boolean tasks

Having established that a single L5PC can solve naturalistic classification tasks, we next asked how far its computational capabilities extend in a more general, abstract setting. While tasks like ’cat’ versus ’dog’ classification are ecologically meaningful, they correspond to specific decision boundaries in a high-dimensional space of spike trains. To systematically probe the neuron’s computational capability, we considered an abstract scenario using Boolean functions.

To illustrate this, consider a scenario in which sensory inputs are first compressed into a small set (e.g., up to 10) of binary features (Fig. S4). Under this representation, each input, such as an image, is mapped to a 10-bit vector, and the neuron is required to spike selectively for a subset of these vectors (e.g., those corresponding to “cats”). With 10 features, the input space contains 1024 distinct patterns, and any assignment of labels over this space defines a Boolean function over the input variables. Thus, each of the original sensory discrimination tasks can be viewed as a particular instance of a Boolean classification problem among the vast space of all possible ones.

This abstraction allows us to move beyond specific ecological tasks to a systematic exploration of computational capabilities, including highly nonlinear functions such as XOR and parity, which are known to be nonlinearly separable and therefore require rich internal structure to implement. In our framework, each binary vector is translated into a noisy spike train to reflect biological variability, and we test which Boolean functions the neuron can robustly realize under these conditions.

We first focused on the d-parity problem (Torres-Moreno et al., 2002) precisely because it is a representative example of a subset of highly non-linear classification problems in d dimensions. The parity problem is a generalization of the XOR problem, a canonical nonlinearly separable task. In the basic XOR case (d=2), two binary input variables P_0_, P_1_ can take four combinations (00, 01, 10, 11). The target output must be 1 only when exactly one bit is 1 (01 or 10) and 0 otherwise. This task is strictly non-linearly separable and requires a non-linear operation that a perceptron (simple summation plus a threshold) famously cannot solve (Minsky and Papert 2017). In our implementation, each input bit (P_0_, P_1_) was represented by the activity of a distinct group of presynaptic axons. On each trial, we generated random spike trains in each group of axons and added spike-time jitter to mimic biological noise. The neuron’s task was to emit at least one spike in the decision window if and only if the parity was odd (Fig. 3a).

**Figure 3.**
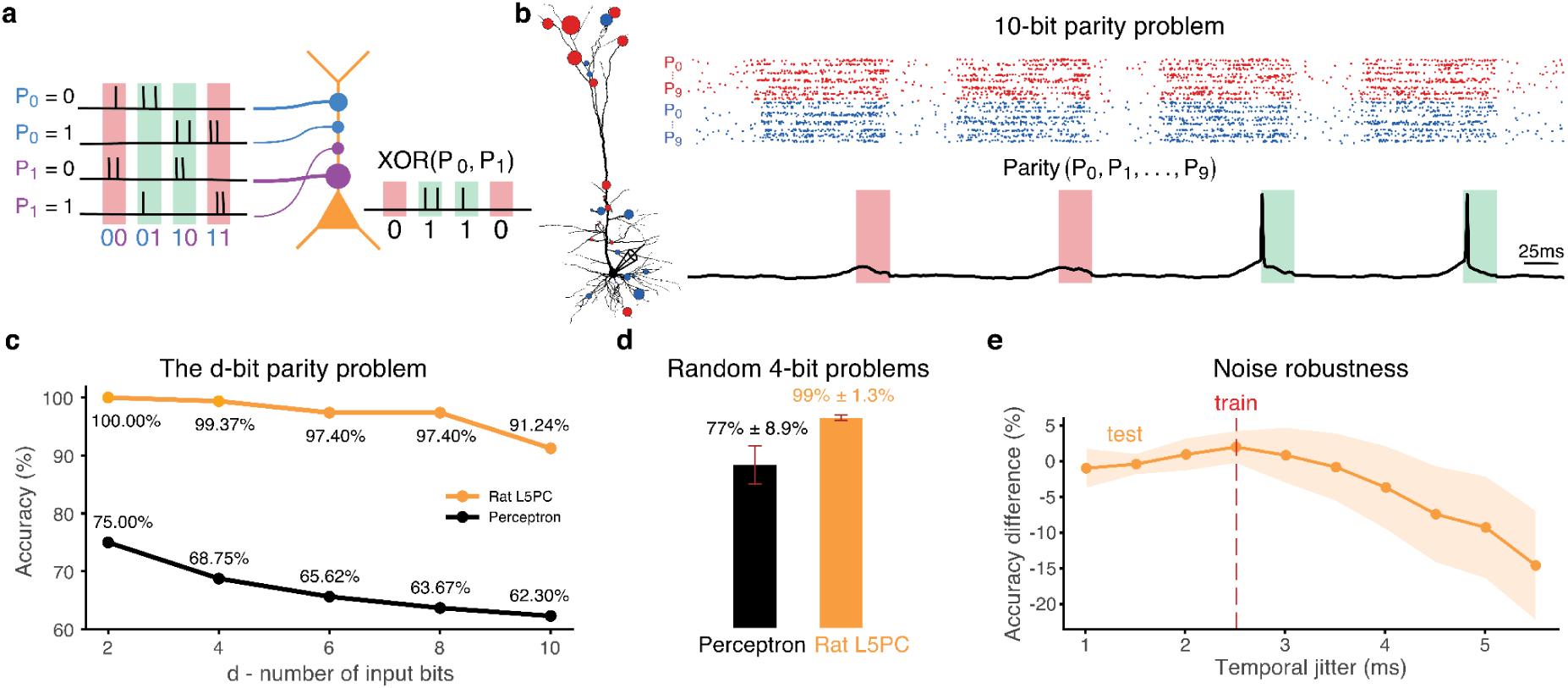
A single L5PC robustly solves arbitrary high-dimensional classification tasks. **a.** Schematic showing a single L5PC challenged with the spatiotemporal XOR problem. The activity of two input axon groups (blue and purple) represents P_0_ and P_1_ variables, respectively. The task is to generate output spikes within a “decision window” of 50 ms only when one variable is active but not when both variables are either silent or active (right; spikes in green windows but not in red windows). **b.** A single L5PC solves the 10-bit parity problem. Left: optimized synaptic weights, obtained by *TwinProp*, for the 10-bit parity problem (the strongest 5% synapses are shown; red - excitatory, blue - inhibitory). Right: Example raster plots of input spikes representing the 10-parity problem (top, subsets of excitatory and inhibitory spikes in red and blue, respectively), and corresponding somatic voltage responses (bottom, in black). Green/Red shaded bars indicate the decision window; for odd-parity patterns, a spike within this window is the correct response (“target”, green window), whereas for even-parity patterns, the correct response is no spike (red shaded decision window). Raster plots span 100 ms and the decision window is 25 ms long. **c.** Accuracy on parity problems, as a function of the problem’s dimension, for L5PC (orange) and perceptron (black). **d.** Performance on random balanced 4-bit problems of L5PC (orange) and perceptron (black). **e.** Robustness of L5PC solution to 4-parity problem to temporal jitter on the test set. Points show mean difference in accuracy with respect to the maximal accuracy (shading, s.d.) as jitter increases; dashed vertical line indicates the jitter in the train set.

To implement the general d-parity problem, we extended the construction discussed above to d input bits (P_0_, P_1_,…, P_d_) by using d disjoint input groups. In the d-parity problem, the output depends on whether the total number of active inputs (# of “1”’s) is odd (for which the output should be 1) or even (for which the output should be 0), see **Methods** and **Fig. S5** for further details. This means that no single input, pair of inputs, or other low-order combination of inputs is predictive on its own, thereby requiring the neuron to nonlinearly integrate information across many inputs simultaneously (Torres-Moreno et al. 2002).

After optimizing synaptic strengths and dendritic locations with *TwinProp* (see respective synaptic strength and location distributions in Fig. S3 A3-F3), the optimized L5PC solved 10-bit parity - a notoriously difficult task whose input space contains 2^10 = 1,024 patterns - with reliable spikes within the decision-window (Fig. 3b). Across dimensionality, accuracy remained near ceiling for d ≤ 8 (pattern length 100 ms, decision window 50 ms) and reached a remarkable accuracy of 91.24% for d = 10 (pattern length 100 ms, decision window 25 ms), consistently surpassing a perceptron baseline, which declined from 75% for d = 2 to 62.3% for d = 10 (Fig. 3c; Torres-Moreno et al., 2002). Thus, a single L5PC can implement computations that require high-order interactions among many inputs, not just linearly separable or low-order conjunctions - a hallmark of a genuine feature binding (see **Discussion**).

Since the d-parity problem constitutes the most computationally demanding non-linear function in d dimensions, our results suggest that the L5PC should theoretically be capable of implementing *any* Boolean function of comparable size. To verify this generality, we next considered random 4-bit Boolean tasks. We focused on 4-bit rules rather than random 10-bit rules because the 2¹⁰ input space becomes prohibitively large for systematic sampling across many distinct decision boundaries, whereas 4-bit tasks permit exhaustive enumeration and controlled class balance while remaining nontrivial. We sampled N = 10 distinct 4-bit rules from the space of balanced Boolean functions, constraining each rule to have the same number of positive and negative outputs across the possible input patterns (see **Methods**). For each rule, we exhaustively presented all input patterns, optimized the synaptic weights of both the L5PC and of a linear perceptron from multiple random initializations, and reported mean test accuracy across rules. The L5PC achieved 99.12 ± 1.34% accuracy across these random tasks, far above the perceptron (76.88 ± 8.86%; Fig. 3d), indicating that a single L5PC can implement a broad range of 4-bit decision boundaries, not only the structured d-parity family.

The resulting solutions were also robust to stochastic temporal perturbations of the inputs. When temporal jitter was added to the input spikes representing the test, accuracy peaked near the jitter used for the training, and declined gracefully thereafter (Fig. 3e), consistent with computation that exploits precise temporal structure rather than purely rate-based pooling (see **Discussion**).

Together, these findings position a single L5 pyramidal cell as a general feature-binding unit, solving both structured (parity) and arbitrary combinatorial classification tasks with temporal precision.

### Distributed NMDA-dependent dendritic computations underlie single neuron solutions

We next examine the dendritic mechanisms that empower the single pyramidal neuron to solve these highly nonlinear complex tasks. During 10-bit parity, membrane voltages across all compartments exhibit rich, arbor-wide activity with decision-locked somatic spikes (Fig. 4a). Thus, correct outputs are accompanied by structured dendritic dynamics rather than purely somatic integration.

**Figure 4.**
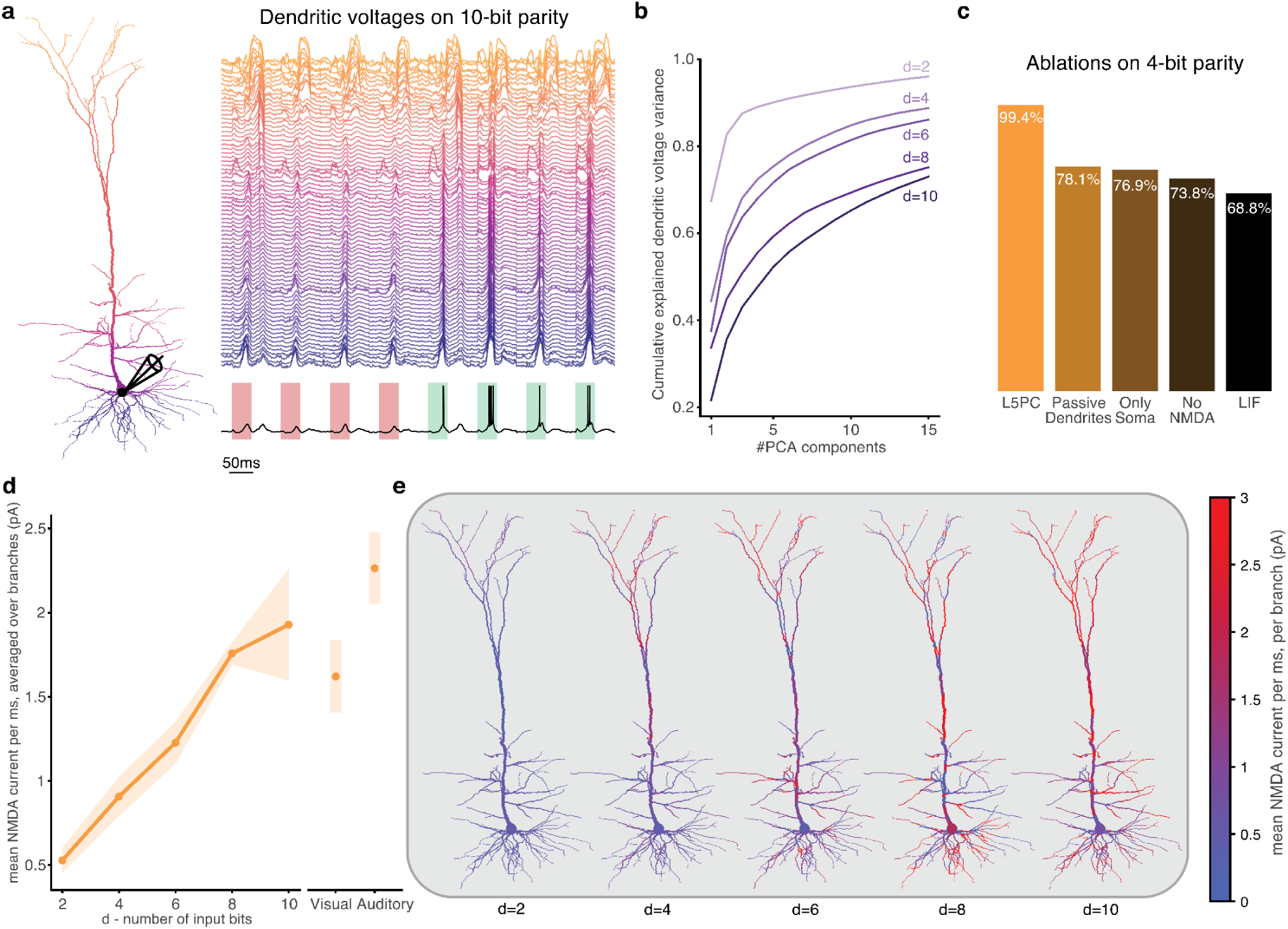
Rich dendritic voltage dynamics and NMDA non-linearities support high-dimensional computations in a single L5PC. **a.** Modeled L5PC (left) and example trial (right) showing membrane voltage from a subset of dendritic compartments of the modeled cell that solved the 10 parity problem (basal - purple, to distal apical tufts - yellow). Somatic voltage is shown in black at bottom; action potentials (in green windows) mark correct stimulus classifications for the problem. **b.** PCA of dendritic voltages: cumulative explained variance as a function of the number of principal components for tasks of increasing dimensionality (d = 2, 4, 6, 8, 10). Higher-dimensional inputs require many more components to explain the variance, indicating richer, higher-dimensional nonlinear dendritic dynamics. **c.** Ablation study on 4-bit parity problem. The fully active L5PC achieves near-perfect accuracy (99.4%). Performance drops with progressively reduced models: passive dendrites (78.1%), soma-only (76.9%), no-NMDA (73.8%), and a LIF neuron (68.8%). **d.** Mean NMDA current per ms increases monotonically with task dimensionality (left, mean ± range across trials), linking more complex computations to stronger recruitment of NMDA-based synaptic conductances. NMDA-recruitment for the naturalistic visual and auditory tasks (Fig. 2) is shown at right, indicating substantial NMDA-dependent dendritic engagement during ecologically relevant computations. **e.** Spatial maps of mean NMDA current per ms across the dendritic tree for d = 2, 4, 6, 8, and 10 parity problems (averaged over samples). Warmer colors denote recruitment of larger NMDA current (color bar shown on right). With increasing d, NMDA engagement becomes more substantial and more spatially widespread, especially in the apical tuft.

To quantify dendritic voltage complexity, we performed principal component analysis, PCA, on dendritic voltages across tasks of increasing dimensionality. The cumulative variance captured by the leading components declines as d - the number of input bits, increases. Namely, with increase in the dimensionality of the problem, more PCA components are required to explain the dendritic voltage patterns (Fig. 4b). This indicates higher-rank, less-shared dendritic activity with increased dimensionality, consistent with many semi-independent dendritic subtrees participating in the respective computation.

We next performed several causal ablation tests to systematically probe which dendritic mechanisms are required to achieve high classification accuracy on the 4-bit parity task. While the intact L5PC performs near-perfectly (99.4%) in solving the task, accuracy drops substantially with any simplification: passive dendrites (voltage-gated channels removed; NMDA retained) - 78.1%; soma-only (morphology collapsed to a single compartment; all channels and NMDA retained) - 76.9%, no NMDA (NMDA removed; voltage-gated channels retained) - 73.8%, and LIF - 68.8% (Fig. 4c). Notably, all three partial simplifications of the detailed neuron model achieve similar sub-optimal performance, well below the intact neuron (see **Discussion**).

To evaluate the contribution of NMDA receptors to the computation, we quantified the mean NMDA current per dendritic segment per millisecond. Mean NMDA current increases monotonically with d (Fig. 4d, left) for the d-parity sets of tasks, indicating that recruitment of NMDA-based synaptic mechanisms scales with task difficulty. The naturalistic visual and auditory tasks (Fig. 2) recruit NMDA currents at levels comparable to high-dimensional parity problems (Fig. 4d, right), indicating that ecologically relevant computations similarly depend on NMDA-mediated dendritic integration. Spatial maps of NMDA activity reveal broader, stronger engagement across the dendritic tree, especially in the apical tuft, as dimensionality of the problem increases (Fig. 4e), suggesting that a larger fraction of the distributed computational compartments of the dendritic tree are utilized.

The analyses above quantify how dendritic voltage structure and NMDA recruitment become richer as task dimensionality increases. To further complement these global scaling results with a fully explicit mechanistic account, we examined the simplest member of the parity family, d = 2, the XOR problem (Fig. S6). Enumerating all four input combinations (P_0_,P_1_ ∈ {0,1}) and inspecting the corresponding excitatory and inhibitory conductances across the dendritic tree, we grouped synapses into two functional subunits, A, corresponding to the apical subtree, and B, corresponding to the basal subtree. We found that subunit B is selectively recruited when P_0_ = 1, P_1_ = 0, whereas subunit A is selectively recruited when P_0_ = 0, P_1_ = 1. Each subunit effectively implements an AND–NOT operation (P_0_ ∧ ¬P_1_ or ¬P_0_ ∧ P_1_) via a specific spatiotemporal arrangement of excitation and inhibition. A nonlinear interaction between these subunits, mediated by a local dendritic calcium spike, combines their outputs such that the soma fires only when exactly one input is active, implementing (P_0_ ∧ ¬P_1_) ∨ (¬P_0_ ∧ P_1_) = P_0_ ⊕ P_1_.

Thus, at the low-dimensional end of the task spectrum, the *TwinProp*-optimized synaptic layout can be reverse-engineered into a compact, interpretable dendritic logic circuit, complementing the distributed, high-rank NMDA-dependent dynamics observed as task dimensionality increases.

In sum, single cortical pyramidal neurons are capable of solving highly nonlinear difficult problems. Implementations of these challenging computations are supported by distributed, NMDA-amplified dendritic integration with high-rank voltage structure. These signatures yield concrete predictions: manipulations that reduce NMDA-based synaptic conductances or restrict dendritic participation (e.g., by shrinking the dendritic tree) will selectively degrade performance, especially for high-dimensional tasks.

## Discussion

A fundamental question in cellular and system neuroscience concerns the computational repertoire of single neurons: what is the set of input–output mappings that a neuron can implement? While it is firmly established that dendritic integration is nonlinear in many neuronal subtypes (Larkum et al. 1999; Schiller et al. 2000; Golding and Spruston 1998; Losonczy and Magee 2006; Llinás and Sugimori 1980), it remains unclear how these mechanisms translate into specific computational benefits for realistic tasks. Here, we operationalize this question by treating the cortical pyramidal cell as a flexible input–output device, asking which target computations can be realized through the optimization of its synaptic weights. Using a differentiable digital DNN twin of the modelled neuron (Fig. 1), we show that a single rat layer-5 pyramidal cell can be trained to solve both naturalistic sensory decision tasks (Fig. 2) and abstract high-dimensional problems from realistic spikes representing the respective inputs (Fig. 3), in both cases far exceeding the capability of perceptron and LIF models (Fig. 2d1, d2, Fig. 3c, d), demonstrating that dendritic mechanisms enable a single neuron to solve complex computational tasks often attributed to multilayer circuits (Fig. 4).

These results are consistent with the view that a pyramidal neuron operates as a network of branches, partially independent submodules with local nonlinearity and global coupling, whose spatially and electrically distributed dendritic structure expands the cell’s representational dimensionality (Polsky et al. 2004; Beniaguev et al. 2021; Poirazi et al. 2003). We link this picture to specific dendritic mechanisms (Fig. 4). PCA of dendritic voltage dynamics shows that, as task dimensionality increases, more components are required, reflecting higher-rank, less-shared dynamics among dendritic arbors and broader arbor engagement (Fig. 4b). In parallel, NMDA currents grow stronger and spread across a wider portion of the dendritic arbor for increasingly challenging parity tasks, especially in apical territories (Fig. 4d, left; Fig. 4e). The same pattern holds for natural image and speech decisions, where NMDA currents are similarly recruited (Fig. 4d, right), suggesting that these naturalistic tasks engage dendritic nonlinearities to a comparable degree. Performance also proved robust to input spike-time jitter, peaking near the training jitter and degrading gracefully (Fig. 3e), consistent with a solution that relies on temporal structure in the input spikes beyond mean firing rates. (Agmon-Snir et al. 1998; Branco et al. 2010; Mainen and Sejnowski 1995). Together, these findings support a mechanism in which multiple, weakly correlated local dendritic events combine to determine whether the neuron fires at decision time, suggesting that a neuron’s optimized synaptic layout can serve as a general feature-binding primitive (Rigotti et al. 2013; Fusi et al. 2016; Johnston et al. 2020; Larkum et al. 1999; Costa and Sjöström 2011; Shai et al. 2015), consistent with in vivo reports of dendritic feature binding (Takahashi et al. 2016; Ranganathan et al. 2018; Zhang et al. 2025; Otor et al. 2022).

Causal ablations close the loop: removing NMDA conductance, silencing voltage-gated dendritic ion channels, or collapsing the dendritic tree to a single compartment each reduces accuracy (Fig. 4c). Notably, removing either NMDA or the dendritic voltage-gated channels yields performance comparable to a soma-only model. Thus, dendritic morphology alone, even with one class of nonlinear mechanism present, does not enhance the neuron’s computational capabilities. Rather, the combination of NMDA-mediated synaptic nonlinearities and voltage-gated dendritic conductances, acting together on the morphology, enriches the computational capabilities of cortical pyramidal neurons. In the simplest parity case (d = 2; Fig. S6), the learned solution can be reverse-engineered into a two-subunit XOR circuit via AND–NOT-like motifs, showing that *TwinProp* can recover mechanistically interpretable synaptic layouts.

These findings generate testable experimental predictions. (i) NMDA antagonists should preferentially impair high-dimensional decisions (Lavzin et al. 2012; Palmer et al. 2014; Grienberger et al. 2014; Smith et al. 2013); (ii) optogenetic inhibition of distal dendritic tufts should preferentially impair complex tasks that rely on widespread dendritic engagement; (iii) voltage imaging should reveal broader, less correlated dendritic activity with more frequent local NMDA events for harder tasks; (iv) learning should correlate with reorganized synaptic placement and increased branch-specific NMDA engagement (Losonczy et al. 2008). A key challenge lies in disentangling single-neuron computation from network-level preprocessing, though emerging methods, including simultaneous imaging of axonal inputs and dendritic responses, and branch-targeted two-photon glutamate imaging and GABA uncaging (Rial Verde et al. 2008; Bazzurro et al. 2022; Aggarwal et al. 2023), may bridge this gap.

Computational capability should scale with morphological and electrophysiological properties (Poirazi and Mel 2001; Aizenbud et al. 2024; Shapira et al. 2025). Large, highly branched arbors with strong active mechanisms, e.g., human L2/3 pyramidal cells, are predicted to support higher-rank solutions than smaller, or more passive cells, e.g., cerebellar granule cells (Otor et al. 2022). Outside the neocortex, Purkinje and CA1/CA3 pyramidal cells offer fertile ground for testing these predictions. *TwinProp* enables, for the first time, converting the “can a single neuron compute X?” question into a quantitative morphology-mechanism-capacity map, applicable across cell types.

*TwinProp* uses digital-twin gradients as an empirical instrument to discover optimized weights, not as a model of biological learning. Yet the solutions map onto known plasticity mechanisms: functional plasticity rules (NMDA- and voltage-dependent) can tune synaptic strengths, while structural plasticity can reroute inputs to appropriate dendritic branches. The high redundancy of cortical representations makes this feasible: a dendrite need not connect to a specific distant axon but can recruit a nearby partner carrying correlated information. The optimized synaptic layouts thus specify concrete hypotheses, which synapses must change, where in the arbor, and by how much, offering a path toward biologically plausible update rules.

Looking ahead, scaling *TwinProp* to multi-cell circuits will clarify the division of labor between dendritic complexity and network architecture (Bast et al. 2024). The framework also enables closed-loop experimental validation: digital twins of specific biological neurons can predict responses to targeted perturbations, directly testing the mechanistic role of dendritic nonlinearities in vivo. In artificial intelligence, these dendritic modules offer a blueprint for bio-inspired architectures that may process spatiotemporal information more efficiently than standard point-neuron networks. Ultimately, integrating single-neuron twins with network-level twins (Wang et al. 2025), emerging volumetric connectomes (Dorkenwald et al. 2024; MICrONS Consortium 2025), and connectome-constrained architectures (Lappalainen et al. 2024; Ito et al. 2026) could yield a multi-scale digital twin of the brain.

## Methods

### Detailed neuron model

We used a detailed compartmental model of a cortical rat layer 5 pyramidal cell (L5PC) as modeled by (Hay et al. 2011). This model contains 12 ion channels per dendritic compartment. Some channels are unevenly distributed across the dendritic arbor. For a complete description of the model, please see STAR Methods in the original paper.

### Detailed synapse models

We used three types of synapses, AMPA, NMDA, and GABAA. AMPA synapses with Double exponential conductance were used in simulations with 𝜏𝑟𝑖𝑠𝑒=0.2ms, 𝜏𝑑𝑒𝑐𝑎𝑦=1.7ms, and typical conductance value of 𝑔𝑚𝑎𝑥=0.4nS. We used the standard NMDA model (Jahr and Stevens 1993), with 𝜏𝑟𝑖𝑠𝑒=0.29ms, 𝜏𝑑𝑒𝑐𝑎𝑦=43ms, 𝛾=0.062mV−1, and typical conductance value of 𝑔𝑚𝑎𝑥=0.3nS. We also used double exponential GABAA synapses with 𝜏𝑟𝑖𝑠𝑒=0.2ms, 𝜏𝑑𝑒𝑐𝑎𝑦=8ms, and typical conductance value of 𝑔𝑚𝑎𝑥=0.7nS. All synaptic parameters were taken from (Markram et al. 2015). Synaptic strengths were later optimized, adhering to biological constraints (see **Synaptic Connectivity and Optimization Constraints** section).

### The *TwinProp* Algorithm

#### *TwinProp* Step 1: digital-twin training

In order to fit a digital-twin DNN model for the simulated neuron, we first followed the study of (Beniaguev et al. 2021), to generate a simulation dataset for the modeled neuron. In each simulation, the modeled neuron was stimulated by random excitatory (AMPA and NMDA) and inhibitory (GABAA) synaptic input with random synaptic weights (both strengths and locations) for a duration of 10s. Each presynaptic spike train was sampled from a Poisson process with a smoothed piecewise constant instantaneous firing rate. The Gaussian smoothing sigma, as well as the time window of constant rate before smoothing, were independently resampled for each 10s simulation from the range of 10 ms to 1000 ms. This was the case, as opposed to choosing a constant firing rate, to create additional temporal variations in the data, in order to increase the applicability of the method to a wide range of potential input regimes. We created a dataset consisting of 50,000 train simulations of 10s each, equivalent to ∼5.7 days of neural data, and 2,000 test simulations of 10s each, equivalent to ∼0.22 days of neural data. Simulations were performed using NEURON software (Carnevale and Hines 2006) and were run in parallel on a CPU cluster.

We then followed (Spieler et al. 2023) to train a DNN with an Expressive Leaky Memory (ELM) architecture on the neuron model simulation datasets. The DNN was fed the same presynaptic spike as the detailed model. The respective DNN was expected to produce a voltage output that matches as closely as possible both the subthreshold and the spiking activity at the soma. In this study, we used an ELM architecture with 1000 memory units and memory time constants ranging from 0.1ms to 300ms, with 3 different random initializations per network. All other parameters were set to be the defaults used originally by (Spieler et al. 2023). Training was executed on NVIDIA GPUs and covered a duration equivalent to approximately 200 days of neural data, corresponding to roughly 35 whole epochs over the entire training dataset. Twin fidelity was assessed using voltage RMSE and spike AUROC on the held-out test set (Fig. S2).

#### *TwinProp* Step 2: synaptic optimization using the twin

With the twin frozen, we optimized synaptic weights to maximize performance on the task. Gradients were computed by automatic differentiation through the twin network. We enforced biological constraints including Dale’s law, contact density limits per axon and dendritic segment, and maximum synaptic conductance (see below **Synaptic Connectivity and Optimization Constraints** section). Optimization was performed using the Adam optimizer with a learning rate in the range [0.001, 0.002]. We employed batched learning (batch size: 32 trials) and minimized the binary cross-entropy loss between the twin’s predicted output spike probability and the target class label. Training ran for 50 epochs. To avoid local minima, we performed multiple restarts (at most 100) with diverse random initial conditions for weight means, variances, and sparsities. All optimization routines were executed on NVIDIA GPUs.

#### *TwinProp* Step 3: validation in the simulation

Optimized synaptic parameters were mapped back to the detailed model. We simulated the full test trials using the detailed model and reported the resulting classification accuracy.

### Synaptic Connectivity and Optimization Constraints

To model realistic afferent connectivity, input axons were not restricted to a single synaptic contact. Instead, we allowed on average 20 synaptic contacts per input axon (excitatory or inhibitory), spatially distributed across the dendritic tree (apical and basal) to maximize nonlinear integration potential. To prevent unrealistic clustering, we enforced a local density constraint ensuring no more than one excitatory synaptic contact and one inhibitory synaptic contact per 1 um of dendritic segment length. Connectivity strictly adhered to Dale’s law: input populations were pre-designated as excitatory or inhibitory, and synaptic weights were constrained to be non-negative. To ensure biological plausibility, synaptic strengths were bounded between 0% and 100% of the typical peak conductance for each synapse type. Maximum conductance values were set to 0.4 nS for AMPA, 0.3 nS for NMDA, and 0.7 nS for GABAA (see **Detailed synapse models** section). The initial synaptic strength distribution was uniform in this range, and during *TwinProp* optimization, weights were clipped to stay within these physiological bounds.

### Baselines

We trained a linear perceptron (Rosenblatt 1958) decoder and a leaky integrate-and-fire (Lapicque 2007) neuron on the same spike inputs and decision rules as the L5PC. The LIF parameters were calibrated to match the passive somatic properties of the L5PC model: membrane time constant 𝜏 = 20 ms, threshold V_th_ = -55 mV, and reset V_reset_ = -80 mV. In addition, we trained fully connected DNN decoders with 2, 5, or 7 layers and 128 units per layer, operating directly on spike representations. The *DNN upper bound* corresponds to the best performance obtained across these architectures.

### Spatiotemporal input encoding protocol

For all tasks, to model realistic presynaptic input, binary input features were encoded by a population of afferent axons using a distributed spatiotemporal protocol. Each binary feature was mapped to a set of N_axons_ presynaptic fibers to ensure robust redundancy. For the visual task, features were mapped uniformly to N_axons_ = 16 fibers each (6,400 axons / 400 features). For the auditory task, features were mapped to N_axons_ = 4 fibers each. For abstract tasks, N_axons_ was scaled such that the total synaptic count remained constant at 8,000 (4,000 excitatory / 4,000 inhibitory). This population utilized a mixed-selectivity coding scheme: a subset of axons encoded the presence of a feature (ON-channels, firing when x_i_ = 1), while the remainder encoded its absence (OFF-channels, firing when x_i_ = 0).

Each axon was assigned a specific target firing time. Across the population, these target times were distributed to tile the full pattern duration (typically 100 ms), ensuring dense temporal coverage. Using a protocol inspired by (Bicknell and Häusser 2021), active axons generated a spike burst centered on their specific target time (mean intra-burst rate of 20 Hz). Spike times were sampled stochastically by applying a temporal Gaussian jitter (σ = 2.5 ms) around the target time.

### Naturalistic sensory tasks

#### Vision

A subset of 1,000 cat and dog images from the AFHQ dataset (Choi et al., 2020) was split 80/20 into training and test sets. Images were preprocessed and passed through a retina → LGN → V1 → V2 pipeline (Riesenhuber and Poggio, 1999) to yield a biologically plausible code of 400 binary features. Spikes were generated from this code via the burst encoding protocol described above, with added temporal noise, resulting in 6,400 excitatory and 6,400 inhibitory axons. The task was to respond to cat images but not to dog images.

#### Audition

A subset of 1,000 speech command waveforms from the SC dataset (Warden, 2018) was split 80/20 into training and test sets. Waveforms were normalized and transformed via a cochlear model (Cramer et al., 2019) and a ventral cochlear nucleus model (Pfeiffer, 1966) into 1,050 binary features reflecting onset/offset/sustained dynamics. Spikes were generated from this code via the same burst-encoding protocol with temporal noise, resulting in 4,200 excitatory and 4,200 inhibitory axons. The task was to respond to the ‘backward’, ‘bird’, ‘dog’, ‘eight’, and ‘follow’ commands, but not to the ‘bed’, ‘cat’, ‘down’, ‘five’, and ‘forward’ commands.

### Abstract Tasks

#### d-parity (generalized XOR)

The d-parity task (Torres-Moreno et al. 2002) serves as a rigorous benchmark for nonlinear capacity, generalizing the exclusive-OR (XOR) problem to d dimensions. The objective is to compute whether the sum of active bits in a d-length binary string is odd. This task is strictly not linearly separable, as changing the state of any single input bit inverts the target label. Spikes were generated from each of bits P_0_, …, P_d-1_ via the burst encoding protocol described above, with added temporal noise, resulting in 4,000 excitatory and 4,000 inhibitory axons. We challenged the neuron with d-parity tasks for d = 2,4,6,8,10.

#### Random Boolean Functions

To test generalizability beyond specific mathematical structures, we evaluated the neuron on random Boolean functions. Truth tables were sampled uniformly from the space of all possible balanced functions over 4 input bits (C(16,8) = 12,870 total functions). While the input spike encoding (burst encoding of bits) remained identical to the parity task, the target output labels were randomized for each task instance, requiring the neuron to learn arbitrary nonlinear mappings.

### Decision readout

Task decisions were read from the somatic spike train within a predefined decision window (see **Results** for the chosen decision window durations). The occurrence of at least one spike within the decision window indicated the positive class. Decoder rules were kept constant across L5PC and baselines to ensure identical information access. Performance was measured as classification accuracy: (TP+TN)/Total, where TP and TN represent true positives and true negatives, respectively.

### Robustness Tests

To evaluate the neuron’s resilience to temporal noise, we perturbed input spike times in the test set with additive Gaussian jitter. We varied the jitter magnitude while holding the training parameters fixed, reporting the degradation in accuracy relative to the baseline performance (no additional jitter).

### Analysis of the XOR mechanism

To dissect the dendritic mechanism solving the XOR (d=2) task (Fig. S6), we analyzed the contributions of distinct morphological compartments. We partitioned the dendritic arbor into two functional subunits: Subunit A (the apical subtree) and Subunit B (the basal subtree). We then simulated the four truth-table conditions (00, 01, 10, 11) and recorded the aggregated excitatory and inhibitory conductances, as well as the local membrane voltage dynamics, separately for each subunit to determine their effective logical operations.

### PCA of Dendritic Voltages

To estimate the effective dimensionality of the dendritic dynamics, we performed Principal Component Analysis (PCA) on the voltage traces. Voltage data was z-scored per compartment to normalize dynamic ranges, then concatenated across time and trials to form a single state matrix. We then computed the principal components and analyzed the cumulative explained variance as a function of the component count for each task difficulty (d).

### Ablations

To isolate the computational contribution of specific morpho-electrical features, we evaluated four model variants: (i) passive dendrites, in which all dendritic voltage-gated channels were removed while NMDA was retained; (ii) soma-only, where morphology was reduced to a single compartment while all channels and synapses were retained; (iii) no-NMDA, in which NMDA conductances were set to zero while voltage-gated channels were retained; and (iv) a LIF control. All variants were trained and tested using identical spike inputs and decision rules.

### NMDA Recruitment

To characterize the involvement of nonlinear mechanisms, we recorded the peak NMDA current at each synapse during task performance. Spatial recruitment maps were generated by averaging these peak currents across trials and mapping the resulting distributions onto the dendritic morphology.

## Acknowledgments

We thank all lab members of the Segev and London Labs for many fruitful discussions and valuable feedback regarding this work. This work was supported by the ONR grant award number N00014-24-1-2055 and grant award number N00014-23-1-2051. M.L. was supported by the ISF grant 1331/23, the NIPI grant 206-22-23, and the BSF grant 2023104. I.S. was supported by the Drahi Family Foundation, the ETH domain for the Blue Brain Project, the Gatsby Charitable Foundation, the NIH grant agreement 1RM1NS132981-01, and by the David and Inez Myers Foundation.

## Author contributions

I.A., conceptualization, methodology, investigation, visualization, software, validation, data curation, writing – original draft; D.B., conceptualization, methodology, writing – review & editing; N.P., investigation, visualization, software; I.S. and M.L., conceptualization, methodology, writing – review & editing, supervision, resources, funding acquisition.

## Code availability

The neuron simulation, digital twin training, neuron utilization via *TwinProp*, as well as code for generating the task datasets (natural and abstract) will be publicly available on GitHub (https://github.com/ido4848/TwinProp) upon publication.

## Data availability

The neuron morphologies and neuron models throughout the whole study will be publicly available on GitHub (https://github.com/ido4848/TwinProp) upon publication.

---

**Figure S1.**
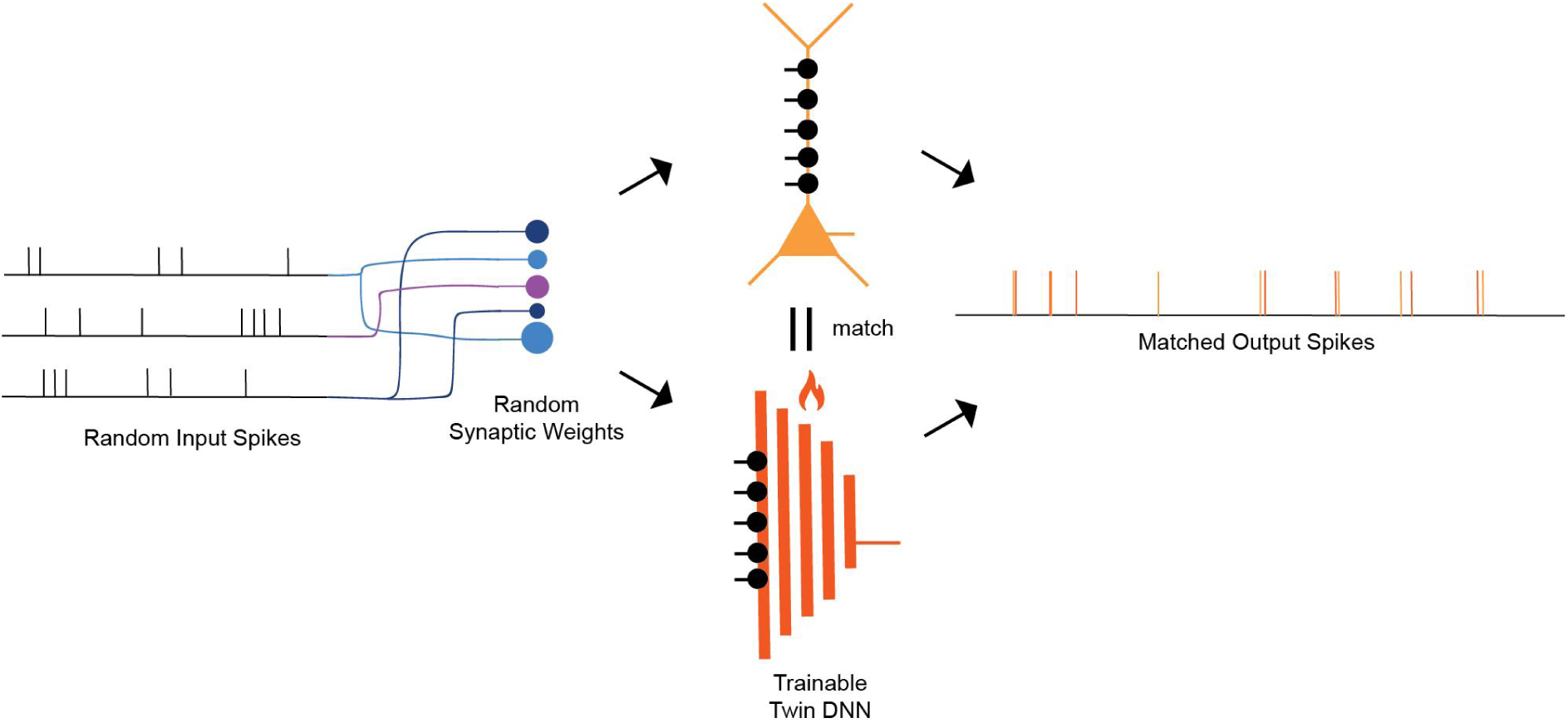
Schematic illustration of the digital twin training phase (*TwinProp* Step 1). To create a differentiable digital twin for the realistic detailed L5PC model, a Deep Neural Network (DNN, bottom center) is trained to mimic the detailed model’s input/output relationship. Both the detailed model (top center) and the DNN are driven by the same large dataset of random input spike trains projected via random synaptic weights (strengths and dendritic locations, left). The trainable parameters of the twin DNN are optimized via gradient descent to match the output spike trains and somatic voltage of the detailed model as closely as possible (right). Once trained, this DNN serves as a frozen, differentiable “digital twin” for subsequent synaptic optimization.

**Figure S2.**
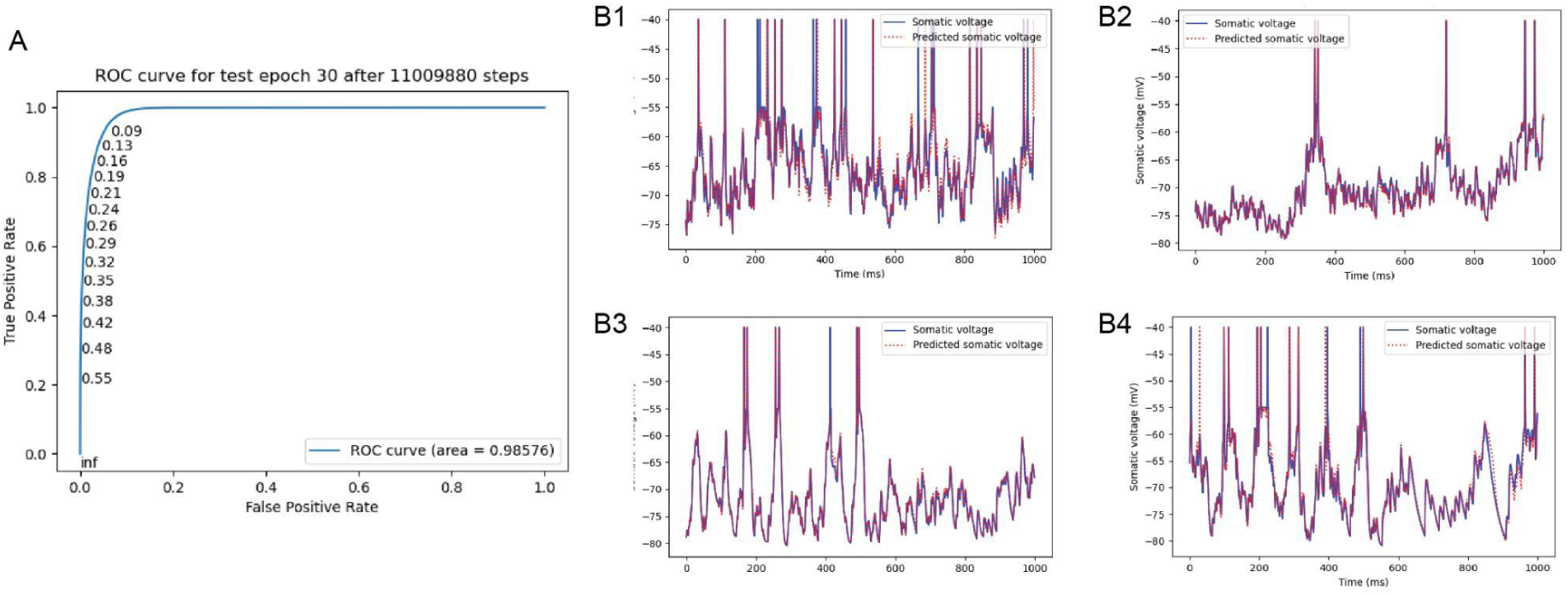
Validation of the digital twin DNN’s predictive accuracy on held-out data. The performance of the trained digital twin DNN was evaluated on a held-out test set of simulations not seen during training. **A.** Receiver Operating Characteristic (ROC) curve illustrating the DNN’s ability to predict the precise timing of somatic spikes generated by the detailed model. The high area under the curve (AUC = 0.98576) indicates excellent spike prediction fidelity. **B1-B4.** Representative examples of somatic voltage traces from the detailed model (solid blue lines) overlaid with the predictions from the digital twin DNN (dashed red lines) given the same inputs. The DNN accurately captures both subthreshold membrane potential fluctuations and spike generation timings across diverse input patterns.

**Figure S3.**
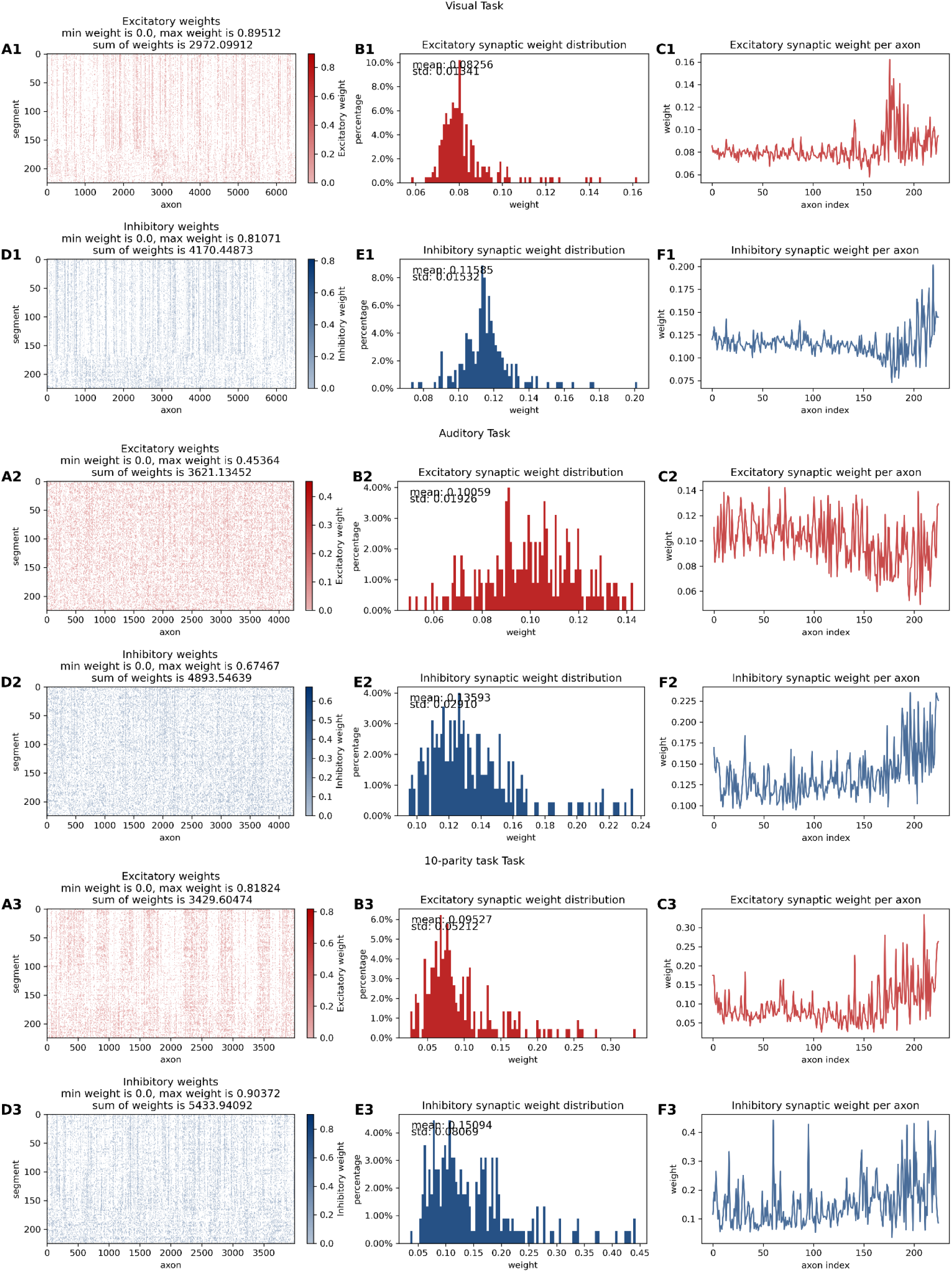
Post-optimization synaptic weight distributions and spatial connectivity across different tasks. Analysis of the synaptic strengths and dendritic locations resulting from *TwinProp* optimization for the Visual classification task (top rows, **A1-F1**), the Auditory classification task (middle rows, **A2-F2**), and the 10-bit parity task (bottom rows, **A3-F3**). For each task, excitatory (red) and inhibitory (blue) synapses are analyzed separately. **A, D.** Visualization of the optimized synaptic weight matrices. Each point represents a synaptic connection between a presynaptic axon (x-axis) and a postsynaptic dendritic segment (y-axis), with color intensity indicating the normalized conductance weight. These matrices reveal the fine-grained “connectome” of the learned solution, highlighting the sparsity and specific dendritic targeting of the inputs. **B, E.** Histograms showing the distribution of optimized synaptic conductance weights (normalized to the maximum physiological conductance for each synapse type). Weights are bounded between 0 and the maximum biologically plausible value, showing that *TwinProp* finds solutions within physiological constraints. **C, F.** Average synaptic weight per presynaptic axon index, showing heterogeneous recruitment of different input fibers.

**Figure S4.**
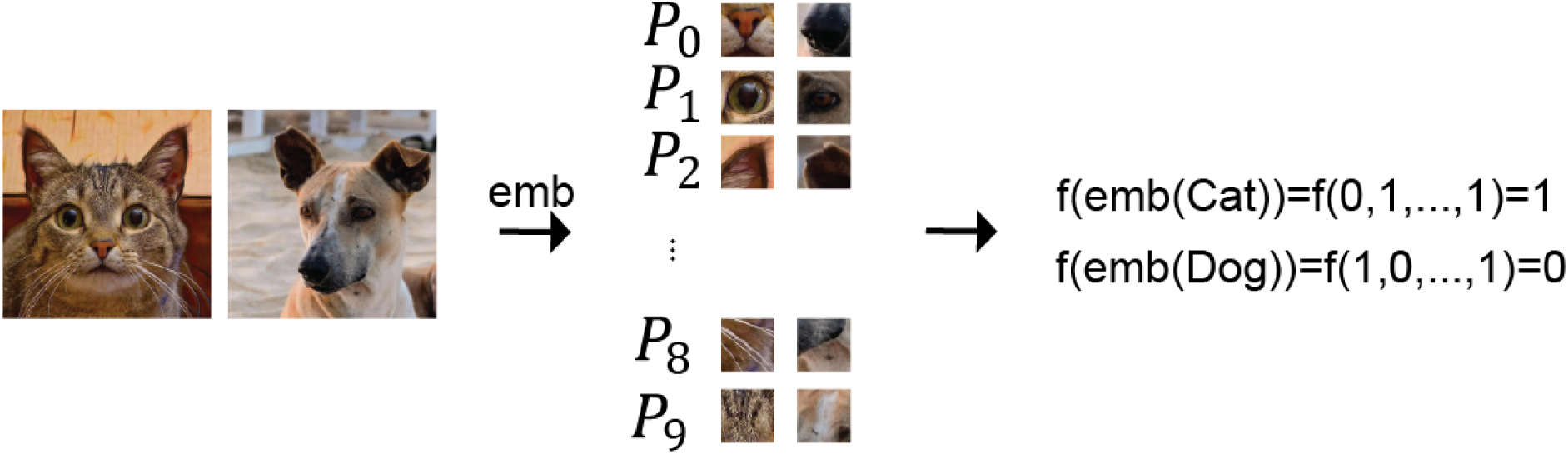
A conceptual illustration of a high-dimensional binary classification task. Complex inputs (e.g., images of a cat or dog) are embedded (”emb”) into a set of discrete features (P_0_, …, P_9_). These features correspond to binary input patterns presented to the neuron function, f. The neuron is tasked with classifying the input correctly (e.g., outputting a spike ’1’ for a cat and remaining silent ’0’ for a dog) based on these high-dimensional binary vectors.

**Figure S5.**
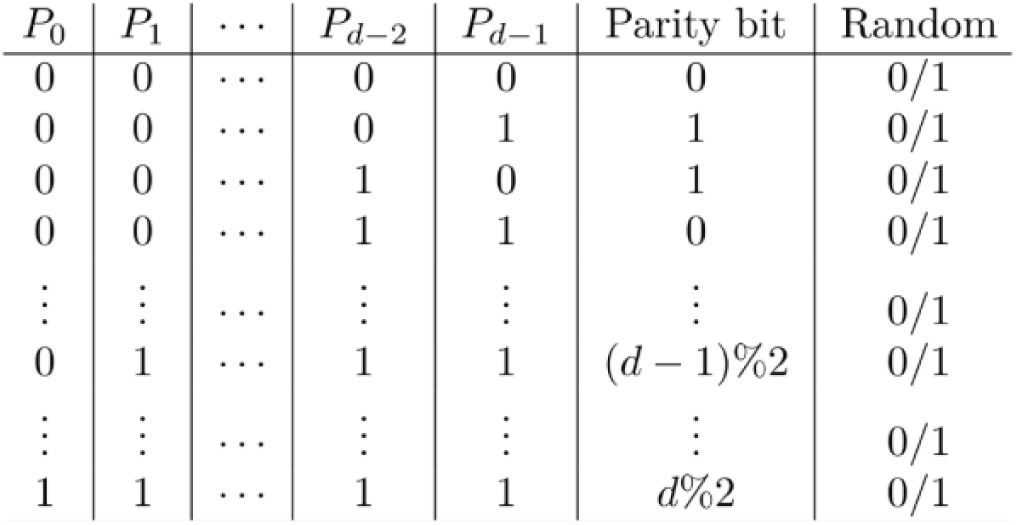
The d-parity task and arbitrary boolean functions. Truth table illustrating the d-parity problem and its extension to arbitrary boolean functions. For the d-parity task, the target output is determined by the sum of the d binary input bits (P_0_, …, P_d-1_): the target is 1 if the sum is odd (Parity bit = 1) and 0 if the sum is even. This represents a rigorous generalization of the non-linearly separable XOR task to higher dimensions. The rightmost column (”Random”) illustrates the more general case of arbitrary boolean functions, in which the target output for each input pattern is assigned an independent binary value (0 or 1).

**Figure S6.**
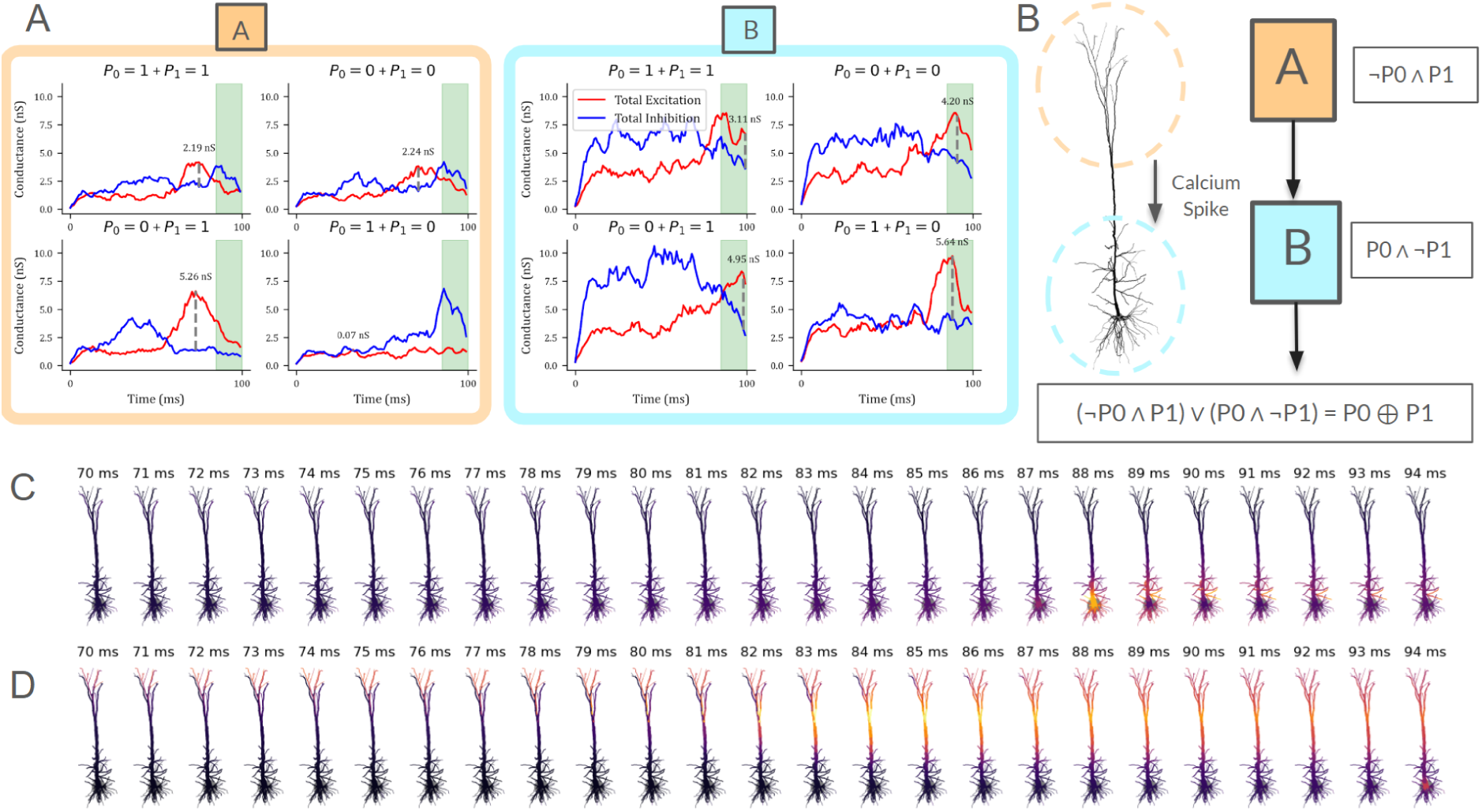
Reverse-engineering the dendritic mechanism of the XOR (d=2) solution. **A.** Analysis of total excitatory (red) and inhibitory (blue) conductances within two functionally distinct dendritic subunits during the four possible input combinations of the XOR task (P_0_, P_1_ ∈ {0,1}). The shaded green region indicates the decision window. Left (Orange Box, Subunit A): This subunit, located in the apical tuft, is selectively active (high excitation, low inhibition) only when P_0_=0 and P_1_=1. Right (Blue Box, Subunit B): This subunit, located in the basal dendrites, is selectively active (high excitation, low inhibition) only when P_0_=1 and P_1_=0. **B.** Schematic summary of the learned solution. The neuron segregates the XOR problem into two linearly separable AND-NOT operations mapped to distinct morphological compartments. The apical tuft (Subunit A) computes ¬P_0_ ∧ P_1_, while the basal tree (Subunit B) computes P_0_ ∧ ¬P_1_. These local signals are integrated (potentially via a calcium spike mechanism) to drive somatic firing if either condition is met, effectively implementing the logical OR of the two branches: (¬P_0_ ∧ P_1_) ∨ (P_0_ ∧ ¬P_1_) = P_0_ ⊕ P_1_. **C, D.** Spatiotemporal voltage heatmaps of the dendritic arbor during the two “odd” input cases. **C.** For input 10 (P_0_=1, P_1_=0), voltage depolarization is confined primarily to the basal dendrites (Subunit B). **D.** For input 01 (P_0_=0, P_1_=1), voltage depolarization occurs primarily in the apical tuft (Subunit A), confirming the spatial segregation of the logic operations.

## References

1. Abbott, L. F. 1999. “Lapicque’s Introduction of the Integrate-and-Fire Model Neuron (1907).” Brain Research Bulletin 50 (5-6): 303–304.

2. Aggarwal, Abhi, Rui Liu, Yang Chen, et al. 2023. “Glutamate Indicators with Improved Activation Kinetics and Localization for Imaging Synaptic Transmission.” Nature Methods 20 (6): 925–934.

3. Agmon-Snir, H., C. E. Carr, and J. Rinzel. 1998. “The Role of Dendrites in Auditory Coincidence Detection.” Nature 393 (6682): 268–272.

4. Agrawal, Anamika, and Michael A. Buice. 2025. “Bounds on the Computational Complexity of Neurons due to Dendritic Morphology.” In Neuroscience, No. Biorxiv;2025.07.11.664448v1. BioRxiv, July 17. https://www.biorxiv.org/content/10.1101/2025.07.11.664448v1.

5. Aizenbud, Ido, Daniela Yoeli, David Beniaguev, Christiaan Pj de Kock, Michael London, and Idan Segev. 2024. “What Makes Human Cortical Pyramidal Neurons Functionally Complex.” In bioRxivorg. December 19. 10.1101/2024.12.17.628883.

6. Bast, Arco, Rieke Fruengel, Christiaan P. J. de Kock, and Marcel Oberlaender. 2024. “Network-Neuron Interactions Underlying Sensory Responses of Layer 5 Pyramidal Tract Neurons in Barrel Cortex.” PLoS Computational Biology 20 (4): e1011468.

7. Bazzurro, Virginia, Elena Gatta, Elena Angeli, et al. 2022. “Involvement of GABAA Receptors Containing α6 Subtypes in Antisecretory Factor Activity on Rat Cerebellar Granule Cells Studied by Two-Photon Uncaging.” The European Journal of Neuroscience 56 (5): 4505–4513.

8. Beniaguev, David. 2023. “Single Biological Neurons as Temporally Precise Spatio-Temporal Pattern Recognizers.” In *arXiv [q-bio.NC]*. September 26. arXiv. http://arxiv.org/abs/2309.15090.

9. Beniaguev, David, Idan Segev, and Michael London. 2021. “Single Cortical Neurons as Deep Artificial Neural Networks.” Neuron 109 (17): 2727–2739.e3.

10. Bicknell, Brendan A., and Michael Häusser. 2021. “A Synaptic Learning Rule for Exploiting Nonlinear Dendritic Computation.” Neuron 109 (24): 4001–4017.e10.

11. Branco, Tiago, Beverley A. Clark, and Michael Häusser. 2010. “Dendritic Discrimination of Temporal Input Sequences in Cortical Neurons.” Science (New York, N.Y.) 329 (5999): 1671–1675.

12. Carnevale, Nicholas T., and Michael L. Hines. 2006. The NEURON Book. Cambridge University Press.

13. Choi, Yunjey, Youngjung Uh, Jaejun Yoo, and Jung-Woo Ha. 2020. “StarGAN v2: Diverse Image Synthesis for Multiple Domains.” Paper presented 2020 IEEE/CVF Conference on Computer Vision and Pattern Recognition (CVPR), 2020/6/13-2020/6/19, Seattle, WA, USA. 2020 IEEE/CVF Conference on Computer Vision and Pattern Recognition (CVPR), June. 10.1109/cvpr42600.2020.00821.

14. Citri, Ami, and Robert C. Malenka. 2008. “Synaptic Plasticity: Multiple Forms, Functions, and Mechanisms.” Neuropsychopharmacology: Official Publication of the American College of Neuropsychopharmacology 33 (1): 18–41.

15. Costa, Rui P., and P. Jesper Sjöström. 2011. “One Cell to Rule Them All, and in the Dendrites Bind Them.” Frontiers in Synaptic Neuroscience 3 (September): 5.

16. Cover, Thomas M. 1965. “Geometrical and Statistical Properties of Systems of Linear Inequalities with Applications in Pattern Recognition.” IEEE Transactions on Electronic Computers EC-14 (3): 326–334.

17. Cramer, Benjamin, Yannik Stradmann, Johannes Schemmel, and Friedemann Zenke. 2019. “The Heidelberg Spiking Datasets for the Systematic Evaluation of Spiking Neural Networks.” In *arXiv [cs.NE]*. October 16. arXiv. http://arxiv.org/abs/1910.07407.

18. Deistler, Michael, Kyra L. Kadhim, Matthijs Pals, et al. 2024. “Jaxley: Differentiable Simulation Enables Large-Scale Training of Detailed Biophysical Models of Neural Dynamics.” In bioRxiv. August 21. 10.1101/2024.08.21.608979.

19. Dorkenwald, Sven, Arie Matsliah, Amy R. Sterling, et al. 2024. “Neuronal Wiring Diagram of an Adult Brain.” Nature 634 (8032): 124–138.

20. Fusi, Stefano, Earl K. Miller, and Mattia Rigotti. 2016. “Why Neurons Mix: High Dimensionality for Higher Cognition.” Current Opinion in Neurobiology 37 (April): 66–74.

21. Gidon, Albert, Timothy Adam Zolnik, Pawel Fidzinski, et al. 2020. “Dendritic Action Potentials and Computation in Human Layer 2/3 Cortical Neurons.” *Science (New York*, N.Y.) 367 (6473): 83–87.

22. Golding, Nace L., Nathan P. Staff, and Nelson Spruston. 2002. “Dendritic Spikes as a Mechanism for Cooperative Long-Term Potentiation.” Nature 418 (6895): 326–331.

23. Golding, N. L., and N. Spruston. 1998. “Dendritic Sodium Spikes Are Variable Triggers of Axonal Action Potentials in Hippocampal CA1 Pyramidal Neurons.” Neuron 21 (5): 1189–1200.

24. Grienberger, Christine, Xiaowei Chen, and Arthur Konnerth. 2014. “NMDA Receptor-Dependent Multidendrite Ca(2+) Spikes Required for Hippocampal Burst Firing in Vivo.” Neuron 81 (6): 1274–1281.

25. Hay, Etay, Sean Hill, Felix Schürmann, Henry Markram, and Idan Segev. 2011. “Models of Neocortical Layer 5b Pyramidal Cells Capturing a Wide Range of Dendritic and Perisomatic Active Properties.” PLoS Computational Biology 7 (7): e1002107.

26. Ito, Shinya, Darrell Haufler, Javier Galvan Fraile, et al. 2026. “Deep-Learning-Assisted Simulation of a Cortical Circuit: Integrating Anatomy, Physiology and Function.” In *bioRxiv*. BioRxiv, March 17. 10.64898/2026.03.13.711751.

27. Jahr, C. E., and C. F. Stevens. 1993. “Calcium Permeability of the N-Methyl-D-Aspartate Receptor Channel in Hippocampal Neurons in Culture.” Proceedings of the National Academy of Sciences of the United States of America 90 (24): 11573–11577.

28. Johnston, W. Jeffrey, Stephanie E. Palmer, and David J. Freedman. 2020. “Nonlinear Mixed Selectivity Supports Reliable Neural Computation.” PLoS Computational Biology 16 (2): e1007544.

29. Jones, Ilenna Simone, and Konrad Paul Kording. 2021. “Might a Single Neuron Solve Interesting Machine Learning Problems through Successive Computations on Its Dendritic Tree?” Neural Computation 33 (6): 1554–1571.

30. Khaligh-Razavi, Seyed-Mahdi, and Nikolaus Kriegeskorte. 2014. “Deep Supervised, but Not Unsupervised, Models May Explain IT Cortical Representation.” PLoS Computational Biology 10 (11): e1003915.

31. Koch, C., and I. Segev. 2000. “The Role of Single Neurons in Information Processing.” Nature Neuroscience 3 Suppl (S11): 1171–1177.

32. Lamprecht, Raphael, and Joseph LeDoux. 2004. “Structural Plasticity and Memory.” Nature Reviews. Neuroscience 5 (1): 45–54.

33. Lapicque, Louis. 2007. “Quantitative Investigations of Electrical Nerve Excitation Treated as Polarization. 1907.” Biological Cybernetics 97 (5-6): 341–349.

34. Lappalainen, Janne K., Fabian D. Tschopp, Sridhama Prakhya, et al. 2024. “Connectome-Constrained Networks Predict Neural Activity across the Fly Visual System.” Nature 634 (8036): 1132–1140.

35. Larkum, M. E., J. J. Zhu, and B. Sakmann. 1999. “A New Cellular Mechanism for Coupling Inputs Arriving at Different Cortical Layers.” Nature 398 (6725): 338–341.

36. Lavzin, Maria, Sophia Rapoport, Alon Polsky, Liora Garion, and Jackie Schiller. 2012. “Nonlinear Dendritic Processing Determines Angular Tuning of Barrel Cortex Neurons in Vivo.” Nature 490 (7420): 397–401.

37. LeCun, Yann, Yoshua Bengio, and Geoffrey Hinton. 2015. “Deep Learning.” Nature 521 (7553): 436–444.

38. Llinás, R., and M. Sugimori. 1980. “Electrophysiological Properties of in Vitro Purkinje Cell Dendrites in Mammalian Cerebellar Slices.” The Journal of Physiology 305 (1): 197–213.

39. London, Michael, and Michael Häusser. 2005. “Dendritic Computation.” Annual Review of Neuroscience 28 (1): 503–532.

40. Losonczy, Attila, and Jeffrey C. Magee. 2006. “Integrative Properties of Radial Oblique Dendrites in Hippocampal CA1 Pyramidal Neurons.” Neuron 50 (2): 291–307.

41. Losonczy, Attila, Judit K. Makara, and Jeffrey C. Magee. 2008. “Compartmentalized Dendritic Plasticity and Input Feature Storage in Neurons.” Nature 452 (7186): 436–441.

42. Magee, J. C. 2000. “Dendritic Integration of Excitatory Synaptic Input.” Nature Reviews. Neuroscience 1 (3): 181–190.

43. Mainen, Z. F., and T. J. Sejnowski. 1995. “Reliability of Spike Timing in Neocortical Neurons.” Science (New York, N.Y.) 268 (5216): 1503–1506.

44. Malenka, Robert C., and Mark F. Bear. 2004. “LTP and LTD: An Embarrassment of Riches.” Neuron 44 (1): 5–21.

45. Markram, Henry, Eilif Muller, Srikanth Ramaswamy, et al. 2015. “Reconstruction and Simulation of Neocortical Microcircuitry.” Cell 163 (2): 456–492.

46. McCulloch, Warren, and Walter Pitts. 2021. “A Logical Calculus of the Ideas Immanent in Nervous Activity (1943).” In Ideas That Created the Future. The MIT Press.

47. Mel, Bartlett W. 1992. “NMDA-Based Pattern Discrimination in a Modeled Cortical Neuron.” Neural Computation 4 (4): 502–517.

48. MICrONS Consortium. 2025. “Functional Connectomics Spanning Multiple Areas of Mouse Visual Cortex.” Nature 640 (8058): 435–447.

49. Minsky, Marvin, and Seymour A. Papert. 2017. Perceptrons. The MIT Press. MIT Press.

50. Moldwin, Toviah, and Idan Segev. 2020. “Perceptron Learning and Classification in a Modeled Cortical Pyramidal Cell.” Frontiers in Computational Neuroscience 14 (April): 33.

51. Otor, Yara, Shay Achvat, Nathan Cermak, et al. 2022. “Dynamic Compartmental Computations in Tuft Dendrites of Layer 5 Neurons during Motor Behavior.” Science (New York, N.Y.) 376 (6590): 267–275.

52. Palmer, Lucy M., Adam S. Shai, James E. Reeve, Harry L. Anderson, Ole Paulsen, and Matthew E. Larkum. 2014. “NMDA Spikes Enhance Action Potential Generation during Sensory Input.” Nature Neuroscience 17 (3): 383–390.

53. Pfeiffer, R. R. 1966. “Classification of Response Patterns of Spike Discharges for Units in the Cochlear Nucleus: Tone-Burst Stimulation.” Experimental Brain Research 1 (3): 220–235.

54. Poirazi, Panayiota, Terrence Brannon, and Bartlett W. Mel. 2003. “Pyramidal Neuron as Two-Layer Neural Network.” Neuron 37 (6): 989–999.

55. Poirazi, P., and B. W. Mel. 2001. “Impact of Active Dendrites and Structural Plasticity on the Memory Capacity of Neural Tissue.” Neuron 29 (3): 779–796.

56. Polsky, Alon, Bartlett W. Mel, and Jackie Schiller. 2004. “Computational Subunits in Thin Dendrites of Pyramidal Cells.” Nature Neuroscience 7 (6): 621–627.

57. Rall, W. 1957. “Membrane Time Constant of Motoneurons.” Science (New York, N.Y.) 126 (3271): 454.

58. Rall, W. 1959. “Branching Dendritic Trees and Motoneuron Membrane Resistivity.” Experimental Neurology 1 (5): 491–527.

59. Rall, W. 1960. “Membrane Potential Transients and Membrane Time Constant of Motoneurons.” Experimental Neurology 2 (5): 503–532.

60. Rall, W. 1967. “Distinguishing Theoretical Synaptic Potentials Computed for Different Soma-Dendritic Distributions of Synaptic Input.” Journal of Neurophysiology 30 (5): 1138–1168.

61. Ranganathan, Gayathri N., Pierre F. Apostolides, Mark T. Harnett, Ning-Long Xu, Shaul Druckmann, and Jeffrey C. Magee. 2018. “Active Dendritic Integration and Mixed Neocortical Network Representations during an Adaptive Sensing Behavior.” Nature Neuroscience 21 (11): 1583–1590.

62. Rial Verde, Emiliano M., Leonardo Zayat, Roberto Etchenique, and Rafael Yuste. 2008. “Photorelease of GABA with Visible Light Using an Inorganic Caging Group.” Frontiers in Neural Circuits 2 (August): 2.

63. Riesenhuber, M., and T. Poggio. 1999. “Hierarchical Models of Object Recognition in Cortex.” Nature Neuroscience 2 (11): 1019–1025.

64. Rigotti, Mattia, Omri Barak, Melissa R. Warden, et al. 2013. “The Importance of Mixed Selectivity in Complex Cognitive Tasks.” Nature 497 (7451): 585–590.

65. Rosenblatt, F. 1958. “The Perceptron: A Probabilistic Model for Information Storage and Organization in the Brain.” Psychological Review 65 (6): 386–408.

66. Schiller, J., G. Major, H. J. Koester, and Y. Schiller. 2000. “NMDA Spikes in Basal Dendrites of Cortical Pyramidal Neurons.” Nature 404 (6775): 285–289.

67. Segev, I., and M. London. 2000. “Untangling Dendrites with Quantitative Models.” Science (New York, N.Y.) 290 (5492): 744–750.

68. Segev, I., and W. Rall. 1988. “Computational Study of an Excitable Dendritic Spine.” Journal of Neurophysiology 60 (2): 499–523.

69. Segev, I., and W. Rall. 1998. “Excitable Dendrites and Spines: Earlier Theoretical Insights Elucidate Recent Direct Observations.” Trends in Neurosciences 21 (11): 453–460.

70. Sejnowski, Terrence J. 2020. “The Unreasonable Effectiveness of Deep Learning in Artificial Intelligence.” Proceedings of the National Academy of Sciences of the United States of America 117 (48): 30033–30038.

71. Shai, Adam S., Costas A. Anastassiou, Matthew E. Larkum, and Christof Koch. 2015. “Physiology of Layer 5 Pyramidal Neurons in Mouse Primary Visual Cortex: Coincidence Detection through Bursting.” PLoS Computational Biology 11 (3): e1004090.

72. Shapira, Sapir, Ido Aizenbud, Daniela Yoeli, et al. 2025. “Biophysical and Computational Insights from Modeling Human Cortical Pyramidal Neurons.” Frontiers in Neuroscience 19 (1579715): 1579715.

73. Smith, Spencer L., Ikuko T. Smith, Tiago Branco, and Michael Häusser. 2013. “Dendritic Spikes Enhance Stimulus Selectivity in Cortical Neurons in Vivo.” Nature 503 (7474): 115–120.

74. Spieler, Aaron, Nasim Rahaman, Georg Martius, Bernhard Schölkopf, and Anna Levina. 2023. “The Expressive Leaky Memory Neuron: An Efficient and Expressive Phenomenological Neuron Model Can Solve Long-Horizon Tasks.” In *arXiv [cs.NE]*. June 14. arXiv. http://arxiv.org/abs/2306.16922.

75. Spruston, Nelson. 2008. “Pyramidal Neurons: Dendritic Structure and Synaptic Integration.” Nature Reviews. Neuroscience 9 (3): 206–221.

76. Stuart, G. J., and B. Sakmann. 1994. “Active Propagation of Somatic Action Potentials into Neocortical Pyramidal Cell Dendrites.” Nature 367 (6458): 69–72.

77. Takahashi, Naoya, Thomas G. Oertner, Peter Hegemann, and Matthew E. Larkum. 2016. “Active Cortical Dendrites Modulate Perception.” Science (New York, N.Y.) 354 (6319): 1587–1590.

78. Torres-Moreno, J. Manuel, Julio C. Aguilar, and Mirta B. Gordon. 2002. “The Minimum Number of Errors in the N-Parity and Its Solution with an Incremental Neural Network.” Neural Processing Letters 16 (3): 201–210.

79. Wang, Eric Y., Paul G. Fahey, Zhuokun Ding, et al. 2025. “Foundation Model of Neural Activity Predicts Response to New Stimulus Types.” Nature 640 (8058): 470–477.

80. Warden, Pete. 2018. “Speech Commands: A Dataset for Limited-Vocabulary Speech Recognition.” In *arXiv [cs.CL]*. April 9. arXiv. http://arxiv.org/abs/1804.03209.

81. Yamins, Daniel L. K., and James J. DiCarlo. 2016. “Using Goal-Driven Deep Learning Models to Understand Sensory Cortex.” Nature Neuroscience 19 (3): 356–365.

82. Yamins, Daniel L. K., Ha Hong, Charles F. Cadieu, Ethan A. Solomon, Darren Seibert, and James J. DiCarlo. 2014. “Performance-Optimized Hierarchical Models Predict Neural Responses in Higher Visual Cortex.” Proceedings of the National Academy of Sciences of the United States of America 111 (23): 8619–8624.

83. Zhang, Yuan, Yanhe Liu, Yachuang Hu, et al. 2025. “Dendritic Computation for Rule-Based Flexible Categorization.” In bioRxiv. June 4. 10.1101/2025.06.03.657766.

